# MetaSpread: A cancer growth and metastatic spread simulation program in Python

**DOI:** 10.1101/2024.04.09.588670

**Authors:** Alfredo Hernández-Inostroza, Erida Gjini

**Affiliations:** Department of Biomedical Engineering, Instituto Superior Tecnico, University of Lisbon, Lisbon, Portugal; Center for Computational and Stochastic Mathematics, Instituto Superior Tecnico, University of Lisbon, Lisbon, Portugal

**Keywords:** cancer, growth, metastatic spread, multi-scale dynamics, simulation

## Abstract

We develop and provide MetaSpread, an open-source simulation package and interactive program in Python for tumor growth and metastatic spread, based on a mathematical model by Franssen et al. (2019). This paper proposed a hybrid modeling and computational framework where cellular growth and metastatic spread are described and simulated in a spatially explicit manner, accounting for stochastic individual cell dynamics and deterministic dynamics of abiotic factors. This model incorporates several key processes such as the growth and movement of epithelial and mesenchymal cells, the role of the extracellular matrix, diffusion, haptotaxis, circulation and survival of cancer cells in the vasculature, and seeding and growth in secondary sites. In the software that we develop, these growth and metastatic dynamics are programmed using MESA, a Python Package for Agent-based modeling (Masad & Kazil, 2015).

## Statement of need

Models of tumor growth and metastatic spread are critical for understanding the key underlying biological processes and clinical evolution in patients. Mathematical models can be of different level of detail, computational or theoretical, spatial or non-spatial in nature, and can have several mechanisms explicit or implicitly embedded in them, including interaction with resources, biomechanical signals, cellular competition, mutation and migration (Chaplain, 2020; Franssen et al., 2021; Macnamara et al., 2020; Opasic et al., 2020; Sadhukhan & Mishra, 2022; Waclaw et al., 2015). While theoretical and analytical advances remain crucial in mathematical models of cancer, computational approaches that offer direct simulation platforms for efficient numerical exploration, focused study and hypothesis testing are also very much needed. Here, we contribute to this aspect, by offering an open source simulation framework in Python for spatio-temporal progression of tumor and metastatic spread. We build the simulation framework on a hybrid mathematical model developed by Franssen et al. (2019), extending the previous work (Anderson et al., 2000; Anderson & Chaplain, 1998), in close agreement with empirical data (Newton et al., 2015; Sabeh et al., 2009). This contribution aims to bridge the gap between mathematicians, oncologists, biologists, computer scientists and interested researchers working in the field of cancer metastatic progression, in need of a computational framework for interdisciplinary study and collaboration.

## Cancer growth and spread model

A 2-dimensional multigrid hybrid spatial model of cancer dynamics is developed in Python (see Figure 1 for a snapshot illustration). Here we combine the stochastic individual based dynamics of single cells with deterministic dynamics of the abiotic factors. The algorithm for dynamic progression at each time step is depicted in Figure 2. In the tumor site we consider two different cancer cell phenotypes: epithelial (epithelial-like) and mesenchymal (mesenchymal-like) cells. The epithelial-like (E) cancer cells reproduce at a higher rate, but diffuse more slowly than mesenchymal (M) cells, which reproduce at a lower rate but diffuse more rapidly. Furthermore, epithelial cells cannot break through the vasculature wall alone, as they require the presence of mesenchymal cells to be able to intravasate into normal vessel entry-points. The exception to this are ruptured vessels, that allow for the intravasation of any type of cancer cell. The cellular growth and movement in space is modeled considering 2 partial differential equations, where random (diffusion) and non-random (haptotaxis) movement are implemented. The model includes two additional equations: one for the spatio-temporal dynamics of matrix metalloproteinase 2 (MMP-2), a chemical that favors the spread of cancer cells, and another for the degradation of the extracellular matrix (ECM), which also favors the haptotactic movement of the cancer cells. The dimensionless model, as described by (Franssen et al., 2019) in Appendix A of their paper, corresponds to 4 PDEs, where the key variables reflect local densities of epithelial cells (*c*_*E*_) and mesenchymal cells (*c*_*M*_), and concentrations of MMP2 (*m*) and extracellular matrix (*w*):

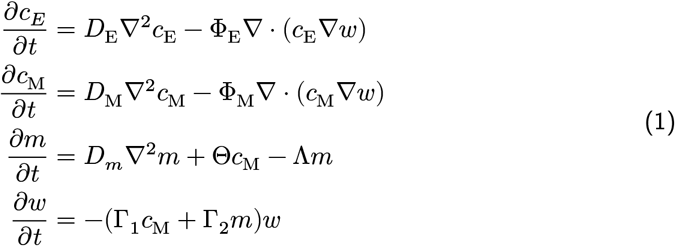

**Figure 1:**
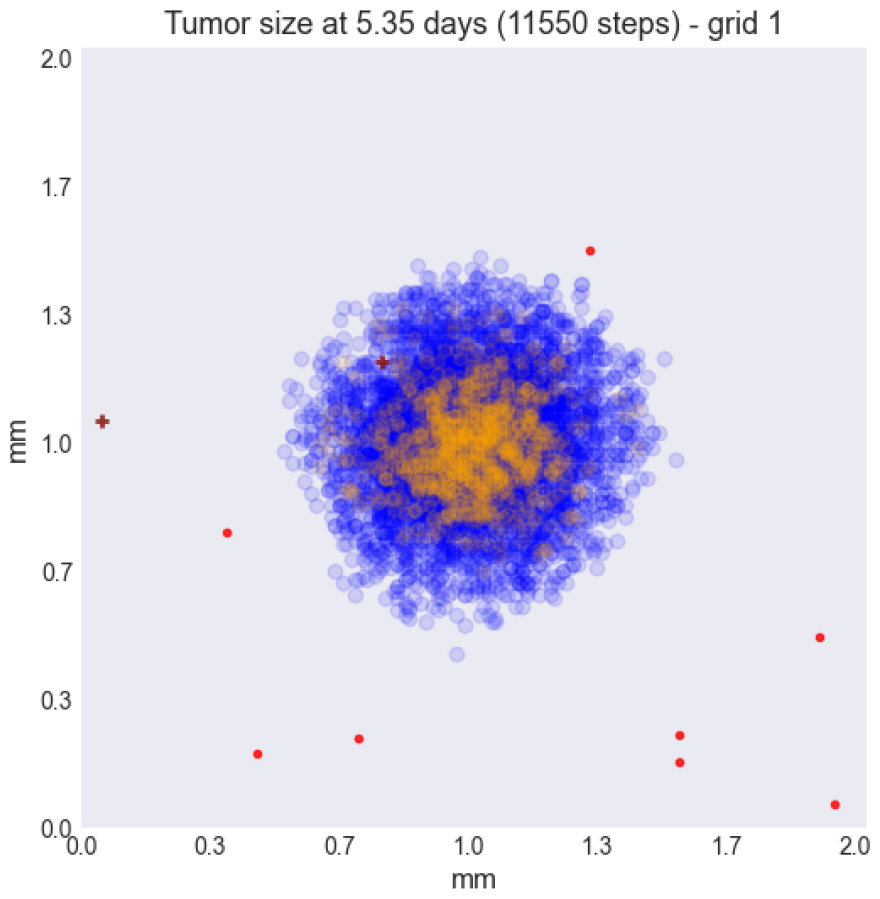
Early snapshot of our simulations for cancer cell spread in the primary tumour (grid 1) after approximately 5 days. Parameters as in Table 1 with initial distribution centered around (1 mm, 1 mm) with radius of about ∼0.1 mm, and total initial size = 388 cells. The blue color denotes mesenchymal cells, the orange color denotes epithelial cells. The intensity of the color represents the number of cells (from 0 to Q = 4) in that particular grid point. The red grid points represent entry-points to the vasculature, with circles intact vessels and crosses representing ruptured vessels.

**Figure 2:**
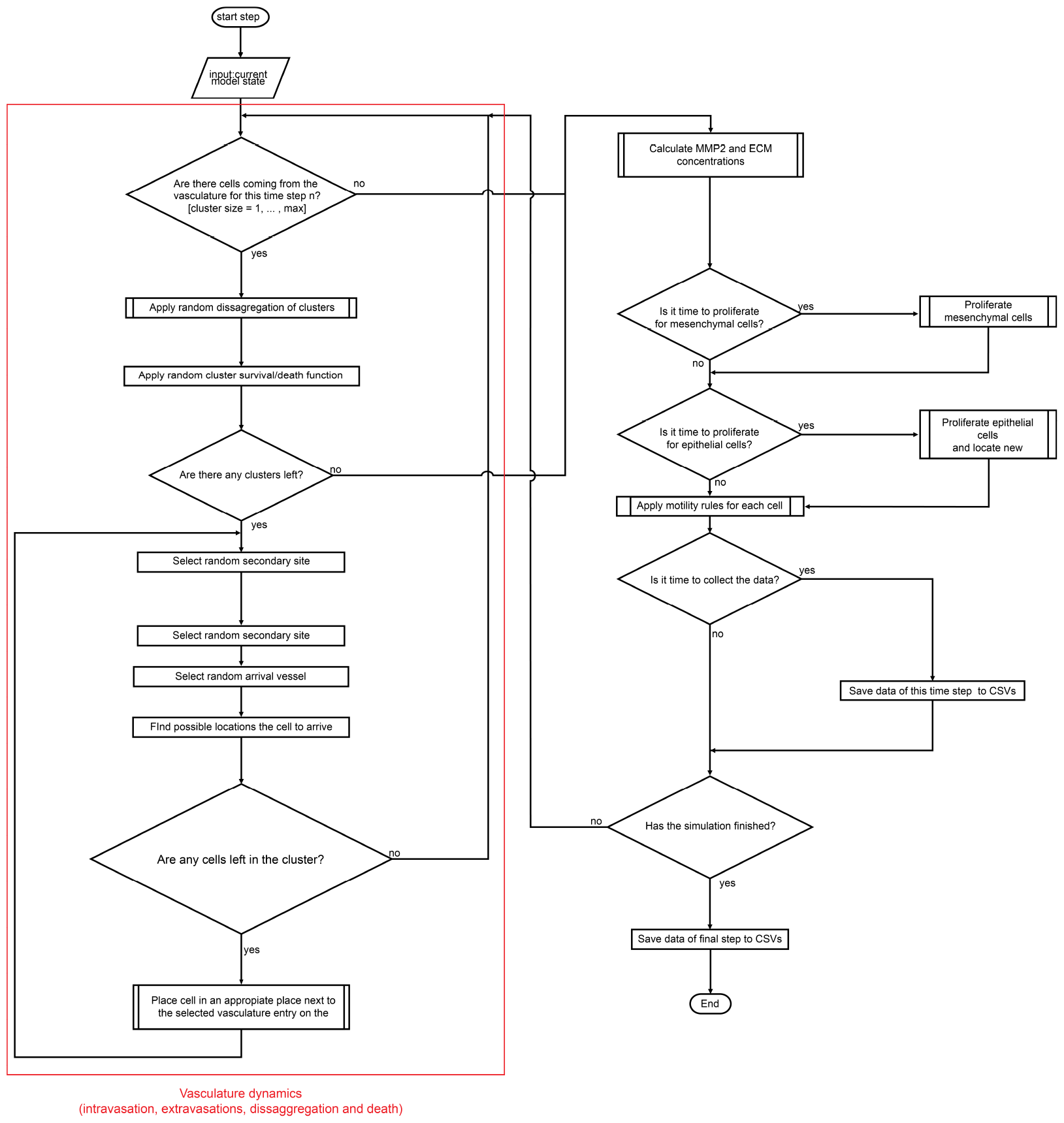
Diagram summarizing the key algorithmic steps of our simulation program in Python, MetaSpread.

For the simulation of the spatio-temporal growth dynamics, and metastatic spread, the system of PDE’s is discretized, and several 2-dimensional grids are established, representing the primary site and the metastatic sites. Discretizing equations for *c*_*E*_ and *c*_*M*_ in space and time, we obtain:

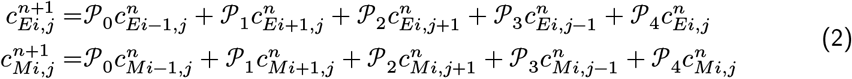

**Table 1:**
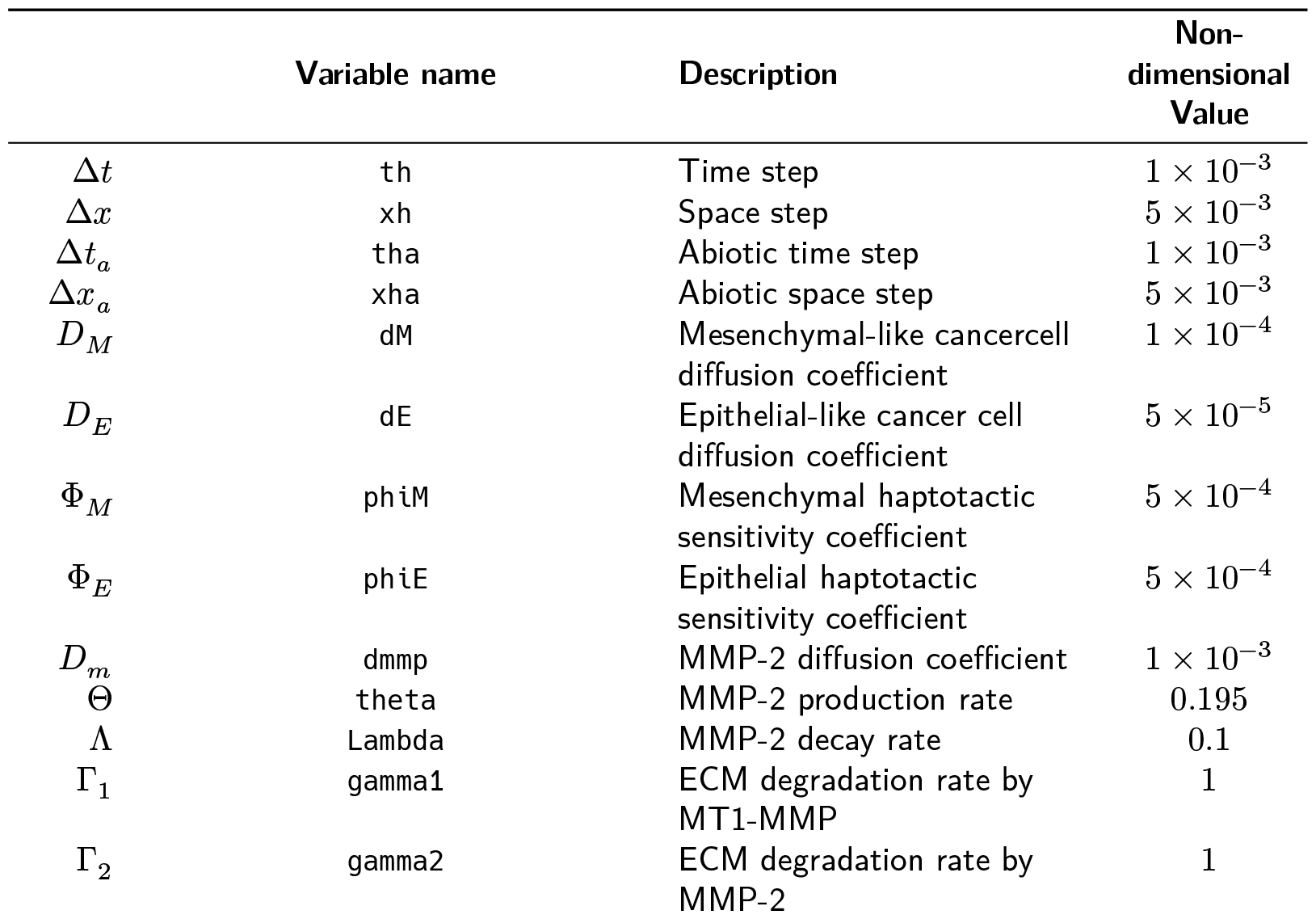
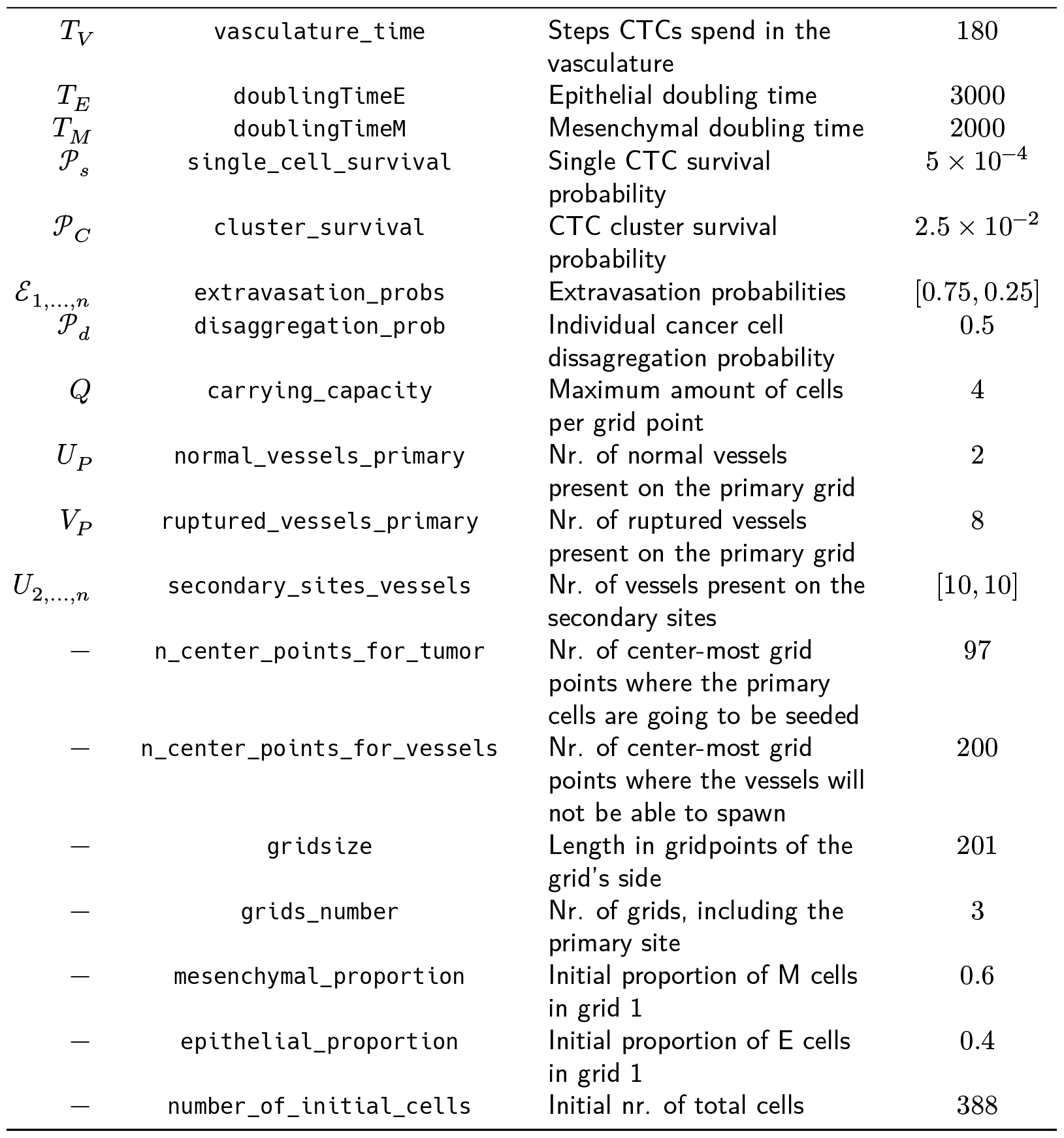
Baseline parameter setup and values used in the computational simulations of MetaSpread. We follow the values estimated and used by Franssen et al. (2019). These parameters are specified in the config file corresponding to each run of the simulation. The non-dimensional values are obtained exactly following (Anderson et al., 2000; Franssen et al., 2019), by scaling time and space with *τ* = *L*^2^/*D* where is a reference diffusion coefficient, an 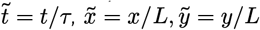, where the original length scale is *L* = 0.2*cm*. With these scalings, the final grid size is 201 x 201.

Where *n* refers to time point, (*i, j*) refers to the spatial grid point (*i, j*), and *𝒫*_0_ to *𝒫*_4_:

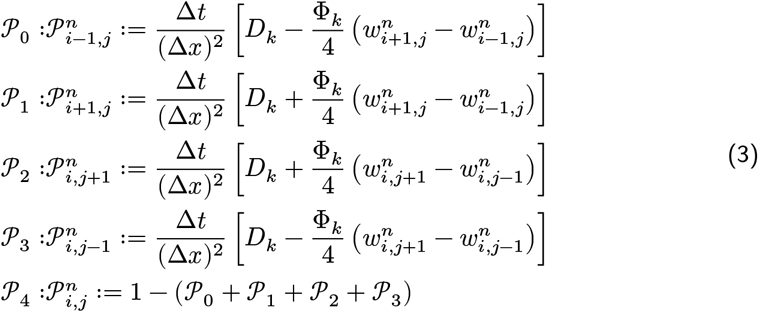

represent the probabilities for a cell to move up, down, left, right, or stay in place, and where *k* = *E, M* can refer to an epithelial-like or mesenchymal-like cell. Each cell on every grid point at location (*x*_*i*_, *y*_*j*_) is modeled as an individual agent, which obeys probability rules for growth and movement. There is a maximal carrying capacity for each grid point given by *Q*, (assumed equal to 4 in (Franssen et al., 2019)), to represent competition for space. There exist a doubling time *T*_*E*_ and *T*_*M*_ for epithelial and mesenchymal cells at which all the cells present in all grids will reproduce, duplicating in place, but never exceeding *Q*.

Only the primary site is seeded with an initial number and distribution of cells. In order for the cells to migrate to another site, they must travel through the vasculature, which they do if they intravasate by one of the several randomly selected points in the grid that represent entrances to the vasculature system. The extravasation to one of the metastatic sites only occurs if they survive, a process that is modeled with net probabilistic rules considering time spent in the vasculature, cluster disaggregation, cell type, and potential biases to different destinations.

For the abiotic factors *m* and *w*, the discretization takes the form (see Appendices in (Franssen et al., 2019)):

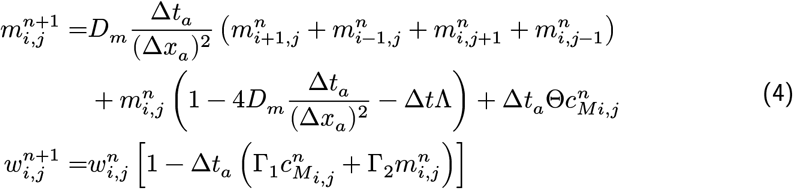

where *i, j* reflect the grid point (*i, j*) and *n* the time-point. In this discretization two different time and spatial steps are used for the cell population (E and M cells) and the abiotic factors (ECM and MMP-2), namely Δ*t* and Δ*x* = Δ*y*, Δ*t*_*a*_ and Δ*x*_*a*_ = Δ*y*_*a*_ respectively.

## Simulation parameters

The biological parameters of the model and the simulation values are summarized in Table 1, tailored to breast cancer progression and early-stage dynamics prior to any treatment and in a pre-angiogenic phase (less than 0.2 cm in diameter). We provide the default values used by (Franssen et al., 2019), as informed by biological and empirical considerations (see also Table 1 and references therein in (Franssen et al., 2019)). The dynamics represent a two-dimensional cross-section of a small avascular tumor and run on a 2-dimensional discrete grid (spatial domain [0, 1] ×[0, 1] corresponding to physical domain of size [0, 0.2] cm ×[0, 0.2]cm), where each grid element corresponds to a spatial unit of dimension (Δ*x*, Δ*y*), and where position *x*_*i*_, *y*_*j*_ corresponds to *i*Δ*x* and *j*Δ*y*. Cancer cells are modeled as discrete agents whose growth and migration dynamics follow probabilistic rules, whereas the abiotic factors MMP2 and extracellular matrix dynamics follow the deterministic PDE evolution, discretized by an explicit five-point central difference discretization scheme together with zero-flux boundary conditions. The challenge of the simulation lies in coupling deterministic and agent-based stochastic dynamics, and in formulating the interface between the primary tumor Grid 1 and the metastatic sites (Grids 2,.. *k*). Each grid shares the same parameters, but there can be biases in connectivity parameters between grids (*ℰ*_*k*_ parameters).

Cell proliferation is implemented locally by generating a new cell when the doubling time is completed, for each cell in each grid point. But if the carrying capacity gets surpassed, then there is no generation of a new cell. The movement of the cells is implemented through the probabilities in Equations 3, which are computed at each time point and for each cell and contain the contribution of the random diffusion process and non-random haptotactic movement. If a cell lands in a grid point that contains a vasculature entry point, it is typically removed from the main grid and added to the vasculature. But there are details regarding the type of cells (E or M) and vasculature entry points (normal or ruptured) further described by (Franssen et al., 2019).

The vasculature is the structure connecting the primary and secondary sites, and it represents a separate compartment in the simulation framework. Single cells or clusters of cells, denominated as circulating tumor cells (CTCs), can enter the vasculature either through a ruptured or normal vessel, and they can remain there for a fixed number of time *T*_*V*_, representing the average time a cancer cell spends in the blood system. Each cell belonging to a cluster in the vasculature can disaggregate with some probability. At the end of the residence time in the vasculature, each cell’s survival is determined randomly with probabilities that are different for single and cluster cells, and the surviving cells are randomly distributed on the secondary sites. To implement this vasculature dynamics in the algorithm, the vasculature is represented as a dictionary where the keys refer to the time-step in which there are clusters ready to extravasate. Intravasation at time *t* corresponds to saving the cells into the dictionary with the associated exit time *t* + *T*_*V*_. It is important to note that this parameter on the configuration file must be in time steps units.

Extravasation rules follow the setup in the original paper (Franssen et al., 2019), ensuring arriving cells do not violate the carrying capacity. Metastatic growth after extravasation follows the same rules as in the original grid.

## Structure of the simulation platform

The program can be run both interactively through the command line, or with explicit user command line arguments.

When run interactively, starting from the main menu, the following possibilities are offered:

▪ **Run a new simulation:** the user can choose the *New Simulation* option to run a new simulation, with the arguments to be specified by the user being the maximal time for the dynamics, and the frequency of saving data (temporal resolution). Any other simulation parameter (see Table 1) will be taken from the *simulation_configs*.*csv* file in the main folder. At the end of the simulation the dynamics of the grids, including agents (cells and vasculature points), the vasculature dynamics and the MMP2 and ECM are saved in a properly identified directory, including a *configs*.*csv* recording the used parameters for this particular simulation. The file *CellsData*.*csv* in this directory will include all the information of all cells and vasculature points in the simulation, for every time step.
  – In addition, in the ECM and MMP2 folders there will be files containing the values of these factors for each time step, not requiring any postprocessing.
  – The vasculature folder will contain several .*json* files with the state of the vasculature at each time step. That is, they will contain a dictionary showing the clusters that were present at each time step. Further information can be extracted by using the **data analysis** option.
  – The folder *Time when grids got populated* will have a file that will simply show the time step for which each grid (primary or secondary site) got populated.
  – When running from the commandline, the user can use `python -m metaspread run max-steps temporal-resolution`. For example, the command `python -m metaspread run 40000 150` would run a simulation for 40000 steps and saving the results every 150 steps.
  – The temporal resolution has to be always less or equal to `vasculature_time`. If not, it will not be possible to see the dynamics of the vasculature correctly, as the cells can intravasate and extravasate without being recorded.
▪ **Load an existing simulation:** The user can select *Load Simulation* from the main menu, and an existing simulation will be loaded, and can be continued for further time steps with the same parameters in its *configs*.*csv* file. The only parameters that the user has to select are the new temporal resolution and the maximum extra steps for the simulation to run. When running from the commandline, the user can use `python -m metaspread load simulation-folder-name additional-steps temporal-resolution`. It is recommended to use the same temporal resolution as used before.
▪ **Post-process data from a simulation:** The generated *CellsData*.*csv* contains the information of every cancer cell at every time step and every grid of the simulation. In order to facilitate the study of the results, we provide the user with several post-processing options: Data analysis, Graphical analysis and Video generation.

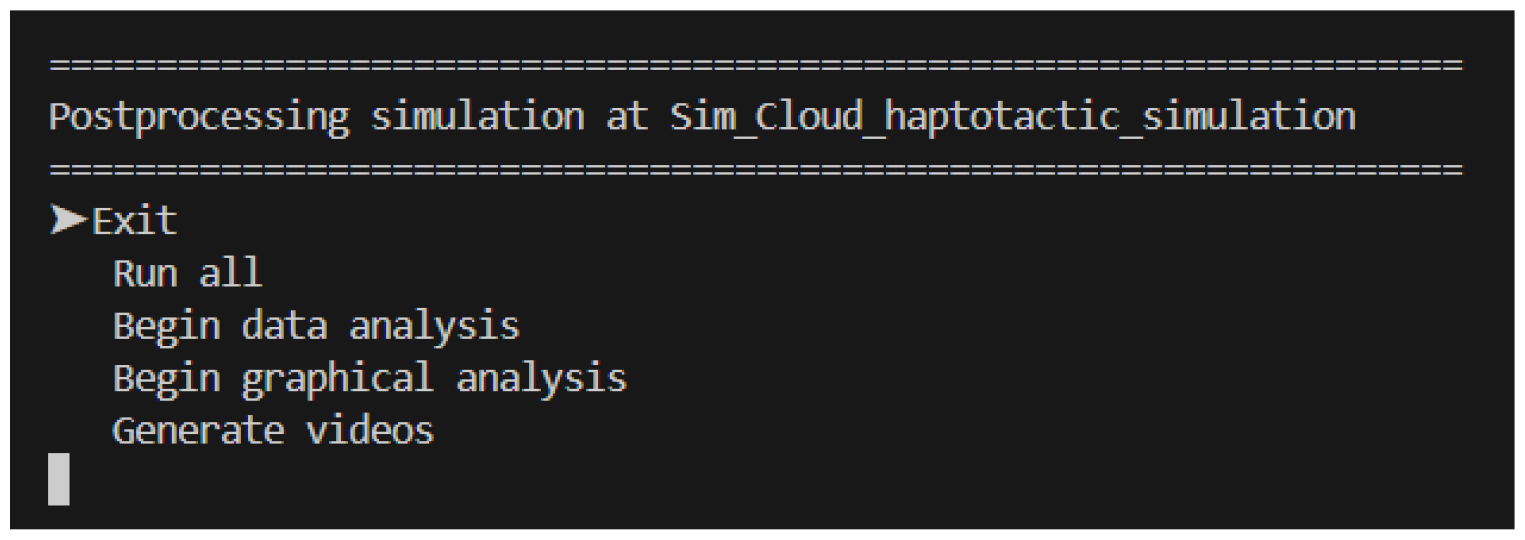
▪ **Data analysis:** several results will be summarized in .*csv* files, such as the vasculature and tumor dynamics.
  – The files that account for total number of cells, Vasculature dynamics (total numbers of CTCs and clusters, cells and phenotypes), and tumor radius (the maximum of all cell distances from the centroid of mass) and diameter (maximum of all cell-to-cell distances) evolution, consist of columns that register the state of a metric in each time step along the simulation. These easily allows plotting graphs of dynamics later on.
  – The tumor growth files for each time point consist of 8 rows: the first 2 rows correspond to x and y coordinates of mesenchymal cells. The second 2 rows correspond to the x and y coordinates of epithelial cells, the next 2 rows correspond to x and y coordinates of regular vasculature points, and the final 2 rows correspond to the coordinates of ruptured vessels. These allow for easily plotting the positions of the agents, and thus, the state of the tumor, at each time step.
  – The histogram files summarize the spatial distribution of cells for each time point. Each file consists of two columns: one for the bins, and one for the frequency. The bins represent the possible number of cells in each grid point, from 0 to *q*, and the frequency the number of grid points that have that amount cells.
  – When running from the commandline, the user can use `python -m metaspread postprocess data simulation-folder-name`
▪ **Graphical analysis:** in order to run this step, it is necessary to run the data analysis option first. When selected, the used will be prompted to introduce the number of figures to describe the snapshot of the dynamics at equally spaced intervals between 0 and the final time of the simulation. Then, plots of the tumor distribution, ECM, MMP-2 for each grid. Furthermore, it will also produce other plots such as the dynamics of the cells in the vasculature, histograms of the cell number distribution over grid points, radius and diameter of the tumor over time, and total size of the tumor in each grid. When running from the commandline, the user can use `python -m metaspread postprocess graphics simulation-folder-name amount-of-figures`.
▪ **Video generation:** The user can choose the Videos option to generate animations from the figures generated in the *graphical analysis* step. When selected, the user will be prompted to introduce the framerate at which the videos should be saved. When running from the commandline, the user can use `python -m metaspread postprocess videos simulation-folder-name frame-rate.`
▪ **Run all:** The user can run all the aforementioned steps in order with this option. When running from the commandline, the user can use `python -m metaspread postprocess all simulation-folder-name amount-of-figures frame-rate`.

## Simulation parameters

The parameters non-dimensional values, as well as their code equivalent name are available in Table 1. Their dimensional values is available in Table 2.

**Table 2:** Additional table with the original values and dimensions of the parameters provided by Franssen et al. (2019).

| Variable | Dimensional Value |
| --- | --- |
| $\Delta t$ | 40 s |
| $\Delta x$ | $1 \times 10^{-3}$ cm |
| $\Delta t_a$ | 40 s |
| $\Delta x_a$ | $1 \times 10^{-3}$ cm |
| $D_M$ | $1 \times 10^{-10}$ cm<sup>2</sup>s<sup>-1</sup> |
| $D_E$ | $5 \times 10^{-11}$ cm<sup>2</sup>s<sup>-1</sup> |
| $\Phi_M$ | $2.6 \times 10^3$ cm<sup>2</sup>M<sup>-1</sup>s<sup>-1</sup> |
| $\Phi_E$ | $2.6 \times 10^3$ cm<sup>2</sup>M<sup>-1</sup>s<sup>-1</sup> |
| $D_m$ | $1 \times 10^{-9}$ cm<sup>2</sup>s<sup>-1</sup> |
| $\Theta$ | $4.875 \times 10^{-6}$ M<sup>-1</sup>s<sup>-1</sup> |
| $\Lambda$ | $2.5 \times 10^{-6}$ s<sup>-1</sup> |
| $\Gamma_1$ | $1 \times 10^{-4}$ s<sup>-1</sup> |
| $\Gamma_2$ | $1 \times 10^{-4}$ M<sup>-1</sup>s<sup>-1</sup> |
| $T_V$ | $7.2 \times 10^3$ s |
| $T_M$ | $1.2 \times 10^5$ s |
| $T_E$ | $8 \times 10^4$ s |

## Simulation output, visualization and analysis

To illustrate the performance and capability of MetaSpread, we provide some figures and visualization of the simulations output. In Figure 3 we show a later snapshot of our simulations for cancer cell spread and ECM and MMP2 evolution. In Figure 4 we show temporal dynamics of summary variables, e.g. total cell counts over time up to 12.78 days, possible to be computed after simulation data post-processing. In movies S1-S2 we show how the simulation platform can be used for studying the biological effect of different perturbations in parameters. These movies illustrate animations of the spatiotemporal evolution of a tumor on the primary site in two cases: (S1) diffusion-dominated and (S2) haptotaxis-dominated cellular movement. The first leads to a regular spatiotemporal pattern of growth, more isotropic and round, the second leads to a more irregular growth over space with cellular protrusions extending in some directions.)

**Figure 3:**
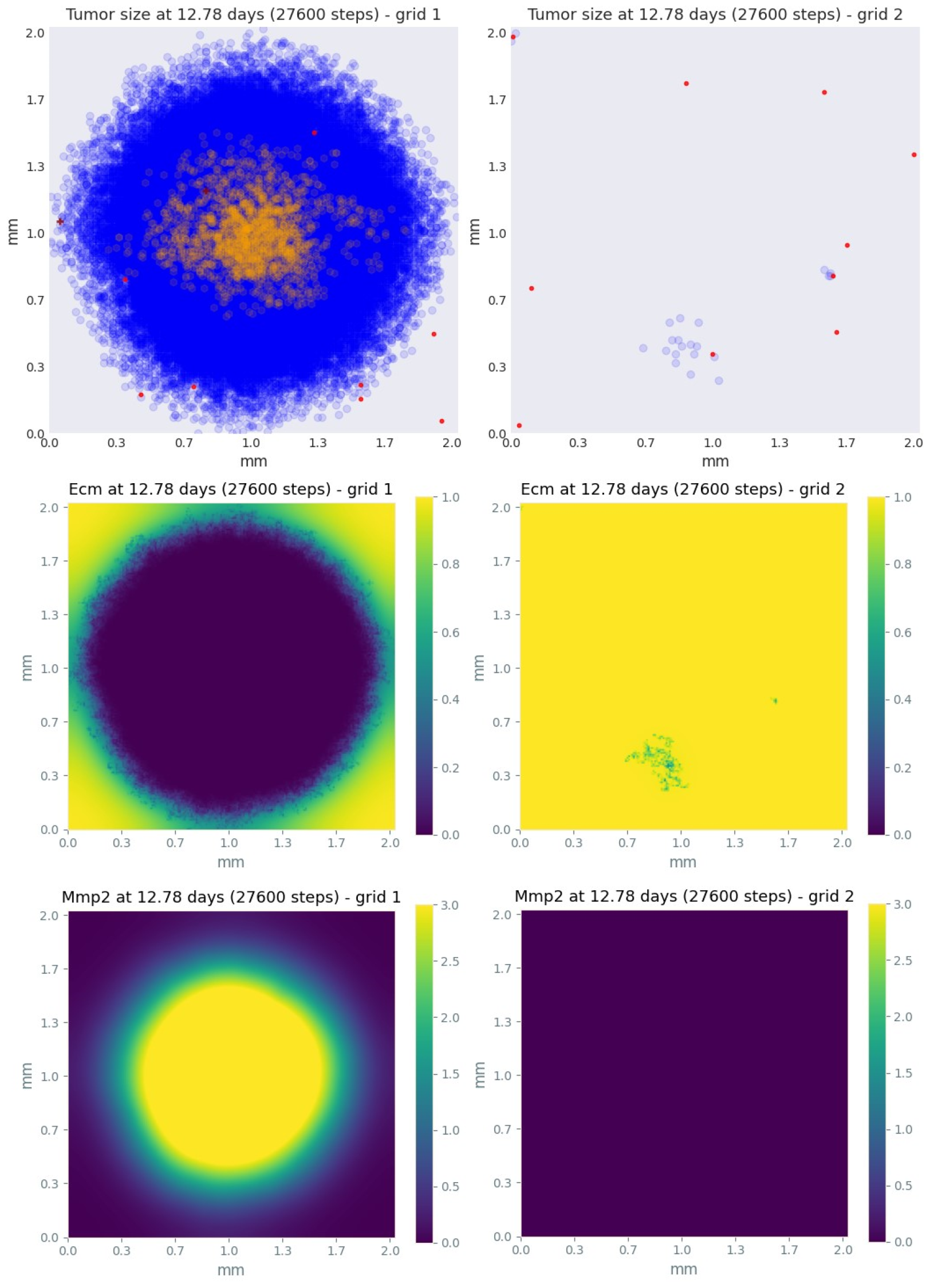
Later snapshot of our simulations for cancer cell spread and ECM and MMP2 evolution in the primary and secondary metastatic site, grid 1 (left) and grid 2 (right) after approximately 12.78 days. Parameters as in Table 1 with initial distribution centered around (1 mm,1 mm) and total initial size = 388 cells. In the top row, the blue color denotes mesenchymal cells, the orange color denotes epithelial cells. The intensity of the color represents the number of cells (from 0 to Q) in that particular grid point. The red grid points represent entry-points to the vasculature, with circles intact vessels and crosses representing ruptured vessels. In the middle row, we plot the corresponding evolution of the density of the extracellular matrix at the same time points. In the last row we plot the spatial distribution of MMP2.

**Figure 4:**
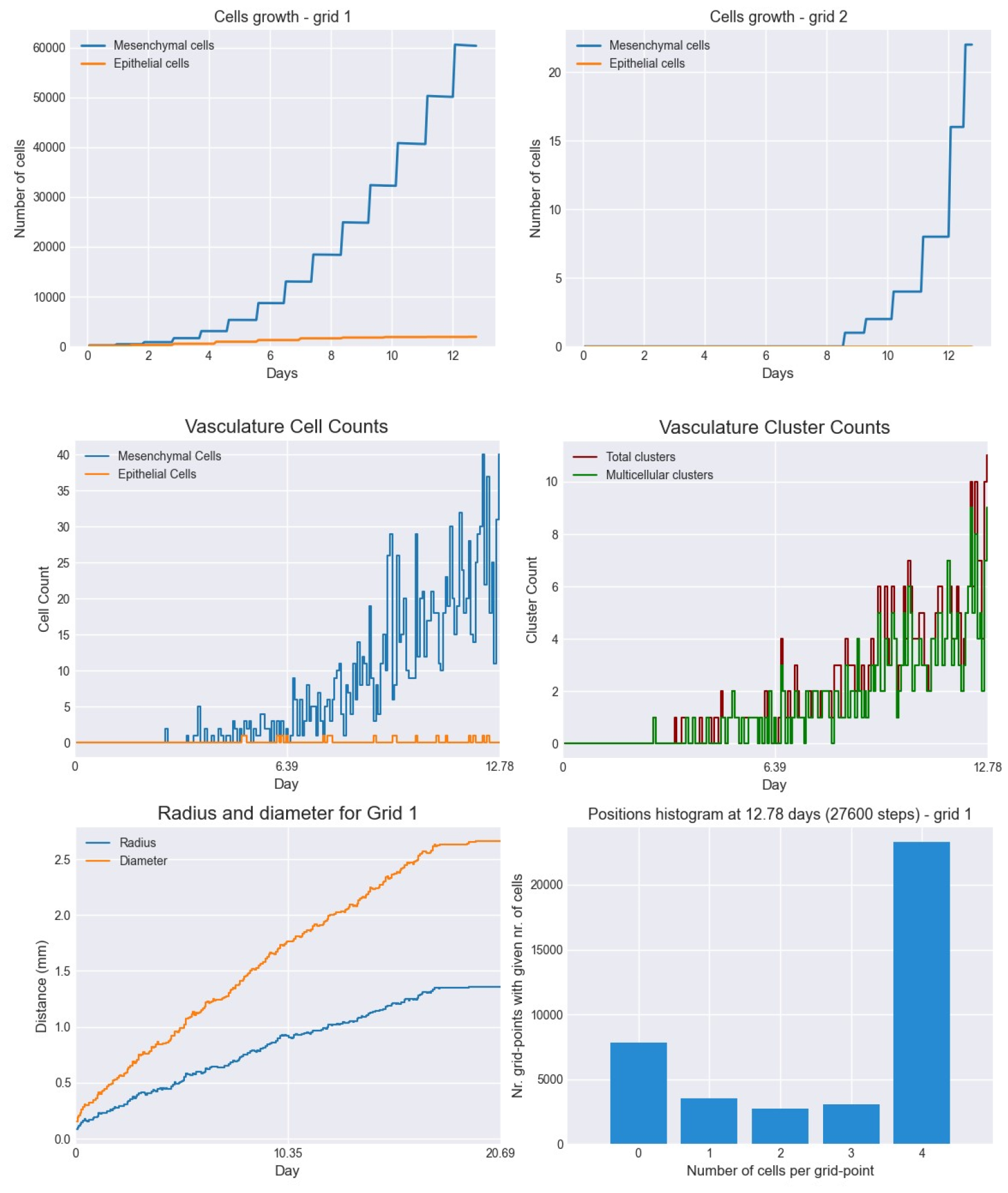
Dynamics of total cell counts over time up to 12.78 days. Top panels: In the primary (left) and secondary (right) tumor grid. Here we illustrate the functionality of the package to yield summaries of the spatiotemporal evolution of the cancer dynamics in the primary and in the metastatic site(s), namely total count of epithelial (E) and mesenchymal (M) cells. Middle panels: Dynamics in the vasculature, showing the amount of E and M cells (left), and the amount clusters (right). Cells can persist as single cells (CTC) or as multicellular clusters. As it can be seen, the majority of cells in the vasculature circulate in the form of clusters (green line) with only a minority being single CTCs (the difference between the red and the green line). Bottom panels: (left) radius and diameter of the spatio-temporal spread Radius is defined as the maximum of all cell distances from the centroid of mass, and diameter as the maximum of all cell-to-cell distances. (Right) distribution histogram of the cells over spatial grid points in the primary grid. The figure is obtained from the simulations corresponding to Figure 3.

## Installation

MetaSpread is available as a official PyPI package. To install, you will need to have PIP installed. Afterwards, run:

~~~
pip install metaspread
~~~

or:

~~~
python -m pip install metaspread
~~~

Manual installation can be done by downloading the repo.

Finally, the program can be run interactiively with:

~~~
python -m metaspread
~~~

Or it can be run purely through command line arguments, as detailed in the Structure of the simulation platform section.

## Outlook

While the model underlying our program (Franssen et al., 2019) is simpler than later models developed for cancer invasion (Chaplain, 2020; Franssen et al., 2021; Macnamara et al., 2020), this simple package enables already deep study of the basic population dynamic processes involved in early tumor dynamics and metastatic growth, and engagement with interesting and important biology (reviewed in Franssen et al. (2019)). A sufficient but not too hard level of complexity makes it a perfect tool for interaction by non-specialists in the mathematical field, medical doctors, and for researchers willing to explore hypotheses with it, perform simulations or extract from it pedagogical value for students and the wider public. There are several directions for extensions of the algorithm and simulation package. These include improving the computational efficiency and speed of the simulation, which now requires about 24 hours for 28.000 time steps, corresponding to about 12 days. Another venue for extension could be including interaction with the immune system, developing explicitly the interaction of the cancer cells with healthy cells, implementing the effect of treatment, for example adaptive therapies (West et al., 2023), mutations and EMT transition. Regarding the metastatic spread, a novelty would be to consider different parameters in different grids, allowing for differential suitability for growth and colonization by arriving cancer cells, which in the current formulation is captured only by the biases in arrival probabilities (Newton et al., 2015). On the computational side the main challenge relies on making the code flexible for parallel computing so that both the spatial and temporal resolution can be increased, and the scope of the phenomena investigated can be expanded, including cell-level heterogeneity (Opasic et al., 2020; Waclaw et al., 2015), and the study of features and variation of many stochastic realizations of the same process. Finally, an interesting improvement would be the addition of an upgraded interface that allows the user to interact with the results in a more user-friendly way, at arbitrary time steps, allowing to probe the tumor dynamics at different degrees of spatial resolution, and selec for enhanced visualization or analysis options.

## Supporting information

Supporting videos are available on the paper branch of the Github repository. Movie S1: Example 1 of spatiotemporal evolution of tumor growth in the primary site (default parameters, diffusion-dominated movement). Movie S2: Example 2 of spatiotemporal evolution of tumor growth in the primary site (parameters with haptotaxis-dominated movement of cells). All the parameters are as default, except for the diffusion coefficients *D*_*M*_ and *D*_*E*_, where in movie 2 they correspond to 1 *⋅* 10^−10^ and 0.5 *⋅* 10^−11^, respectively.

## Acknowledgements

We acknowledge the contributions of Murillo Texeira and Vinicius Schaedler Damin to this project. E. G. acknowledges support by the Portuguese Foundation for Science and Technology (FCT) via CEECIND/03051/2018.

